# Persistent activation of STAT6 in keratinocytes elicits neutrophilic skin inflammation, pruritus and *S. aureus* colonization reminiscent of chronic atopic dermatitis

**DOI:** 10.64898/2026.02.03.703471

**Authors:** Daniel Radtke, E Da Choi, Lisa-Marie Graf, Jonathan Pollock, Daniela Pflaum, Walter Geißdörfer, Stefan Wirtz, Sabine A. Eming, David Voehringer

**Affiliations:** Department of Infection Biology, University Hospital Erlangen and Friedrich-Alexander Universität Erlangen-Nürnberg, Erlangen, Germany; FAU Profile Center Immunomedicine (FAU I-MED), Erlangen, Germany; Microbiology Institute - Clinical Microbiology, Immunology, and Hygiene, University Hospital Erlangen and Friedrich-Alexander Universität Erlangen-Nürnberg, Erlangen, Germany; Department of Medicine 1, Friedrich-Alexander Universität Erlangen-Nürnberg, Erlangen, Germany; Department of Dermatology, University of Cologne, Cologne, Germany; Cluster of Excellence Cellular Stress Responses in Ageing-associated Diseases (CECAD), University of Cologne, Germany; Center for Molecular Medicine Cologne (CMMC), University of Cologne, Germany; Institute of Zoology, Developmental Biology Unit, University of Cologne, Cologne, Germany

## Abstract

Atopic dermatitis (AD) evolves from initial type 2 immunity-driven inflammation to chronic mixed responses by poorly understood mechanisms. To investigate how the prolonged activation of the usually IL-4/IL-13-induced transcription factor STAT6 in keratinocytes impacts on the development and subtype of AD, we generated a new mouse model in which a constitutively active form of STAT6 is selectively expressed in keratinocytes. These K14Cre^+^STAT6^vt/vt^ mice spontaneously developed AD-like skin lesions characterized by *Staphylococcus aureus* colonization, neutrophilic inflammation, and pruritus starting at the age of 12-14 weeks. Treatment with antibiotics mitigated pathology, indicating that it is microbiota-driven. Comparison of human AD gene expression data with the transcriptome of skin biopsies from K14Cre^+^STAT6^vt/vt^ mice revealed features shared with chronic AD, including genes associated with neutrophil and keratinocyte activation. Furthermore, heterozygous K14Cre^+^STAT6^vt/wt^ mice developed a mixed eosinophilic and neutrophilic skin inflammation with exacerbated pathology compared to wild-type controls in an induced model of atopic dermatitis, compatible with chronic AD. These results indicate that persistent STAT6 activity in keratinocytes facilitates *S. aureus* outgrowth on the skin, promotes a type 1-/type 3-biased immune response, and is sufficient to mimic the transition from acute type 2 immunity-to chronic type 1-/type 3-immunity-dominated AD.

## Introduction

Atopic dermatitis (AD) is one of the most common inflammatory skin diseases, with an estimated lifetime prevalence of up to 20%^1, 2^. While disease severity typically peaks in adolescence, chronic disease often persists for years up to a lifetime. Understanding its progression is crucial for effective disease management. AD’s pathogenesis involves complex contributions of the immune system, often transitioning from type 2 immunity to mixed type 1, 2, and 3 responses^3^. AD manifests in diverse clinical presentations, with considerable variability across pediatric and adult forms, as well as region-specific presentations like the Th17-biased Asian form^4, 5, 6^. The drivers of these endotypes remain a critical area of investigation, with ongoing efforts to reconcile underlying mechanisms broadly suggested by the “inside-out” and “outside-in” hypothesis. “Inside out” describes an underlying internal dysregulation of the immune system in the pathogenesis of AD, while “outside in” focuses on external irritants and allergens in driving AD^7^. One striking factor associated with pathogenicity is the prominent colonization of AD skin with *S. aureus*, seen on many AD patients^8^. It can contribute to pathogenicity by production of toxins and other virulence factors but its prominent outgrowth and the resulting dysbiosis might already suffice to enhance AD severity^9, 10, 11^.

In lesional skin of human AD patients, enhanced levels of the alarmin TSLP were reported^12^. In mice, induction of TSLP release by keratinocytes upon topical administration of the vitamin D3 analog MC903 (calcipotriol) promotes an AD-like phenotype and is therefore often used as a model for AD^13, 14, 15^. TSLP activates DCs that then preferentially induce type 2 immunity promoting T cells^12^. In the early phase of AD, the type 2 immunity driving cytokines IL-4 and IL-13 dominate and are sensed by cells via two types of IL-4 receptors. The type I receptor exclusively binds IL-4, while the type II receptor binds IL-4 and IL-13, but both trigger the downstream phosphorylation of STAT6^16, 17, 18^. *In vitro* experiments on keratinocytes show that IL-4 stimulation and STAT6 overexpression reduce the expression of the epidermal differentiation complex, encoding proteins that are relevant to maintain skin-barrier integrity^19,20^. However, it remains unclear, how STAT6-regulated genes in keratinocytes contribute to disease progression in AD ^21^. Selective deletion of IL-4Rα in keratinocytes results in more dendritic epidermal T cells and better wound healing^21^. On the other hand, overexpression of IL-4 or TSLP by - but often not limited to - keratinocytes or their artificial induction triggers type 2 immunity-associated receptors and STAT6 phosphorylation in a variety of cells, leading to acute AD-like pathology ^13, 14, 15, 22, 23^. However, these approaches lack the resolution to define the specific contribution of STAT6-regulated genes in keratinocytes to AD pathogenesis over time.

Here, we utilize a genetic approach to investigate the outcome of persistent STAT6 activity specifically within keratinocytes, mimicking the effector functions induced by IL-4/IL-13 signaling. This cell type-specific resolution of disease pathogenesis, previously unexplored in AD research, aligns with the growing potential of precision medicine to target intracellular pathways in cell type-specific ways^24, 25, 26, 27^. Furthermore, recent identification of gain-of-function STAT6 mutations now classified as a monoallelic human disease^28, 29, 30, 31^ underscores the broader relevance of our findings for understanding skin biology and immune dysregulation.

## Results

### K14Cre^+^STAT6^vt/vt^ mice develop neutrophilic skin inflammation and scratching behavior

To investigate the impact of type 2 immunity-driven keratinocyte activation, we generated K14Cre^+^STAT6^vt/vt^ mice which express a constitutively active mutated form of STAT6 (STAT6vt) under the control of the murine K14 promoter, which is predominantly active in basal epidermal keratinocytes^32^. The STAT6vt mutant has previously been successfully applied to study persistent STAT6 activity in lymphocytes by transgenic expression under control of the human CD2 promoter^33^ or in other cell types using various Cre lines^34, 35, 36^. As expected, phosphorylated STAT6 was observed in ear and chest skin but not in the spleen of K14Cre^+^STAT6^vt/vt^ mice (**Fig. 1A**). At the age of 11–14 weeks, these mice spontaneously developed skin inflammation on the ear and neck region (**Fig. 1B**), also evidenced by quantification of ear swelling (**Fig. 1C**) and loss of skin-barrier integrity indicated by increased transepidermal water loss (TEWL; **Fig. 1D**). At the age of 15–17 weeks, scratching behavior emerged (**Fig. 1E**), accompanied by reduced weight gain (**Fig. 1F**) and development of skin lesions. Increased IgE levels in the serum of 11–14-week-old mice indicated a type 2 immunity-associated antibody response (**Fig. 1G**). To investigate whether IgE was required for skin pathology, we generated bone marrow chimeras by irradiating K14Cre^+^STAT6^vt/vt^ mice and reconstituting them with IgE-deficient (IgEKO) bone marrow^37^. However, IgE did not mediate the skin phenotype, as IgE-deficient K14Cre^+^STAT6^vt/vt^ mice still exhibited ear swelling and barrier leakage (**Suppl. Fig. 1A-B**) ^37^. Analysis of blood leukocytes over time revealed increased neutrophils and Ly6C^+^CD11b^+^ cells at 14 weeks of age, while eosinophils, basophils, CD4^+^ T cells, and CD8^+^ T cells remained unchanged (**Fig. 1H, gating in Suppl. Fig. 2A**). In inflamed ear skin, we observed a significant increase in CD45^+^ cells compared to non-inflamed ears of WT mice (**Fig. 1I**). Among these infiltrating cells, granulocytes were strongly enriched, with pronounced neutrophilia but only modest increases in basophils and eosinophils relative to ears of WT mice (**Fig. 1J-K, gating in Suppl. Fig. 2B**). Thus, persistent STAT6 activity in keratinocytes induces skin inflammation, loss of barrier integrity, and scratching behavior, accompanied by type 3 immunity-associated neutrophilia.

**Figure 1.**
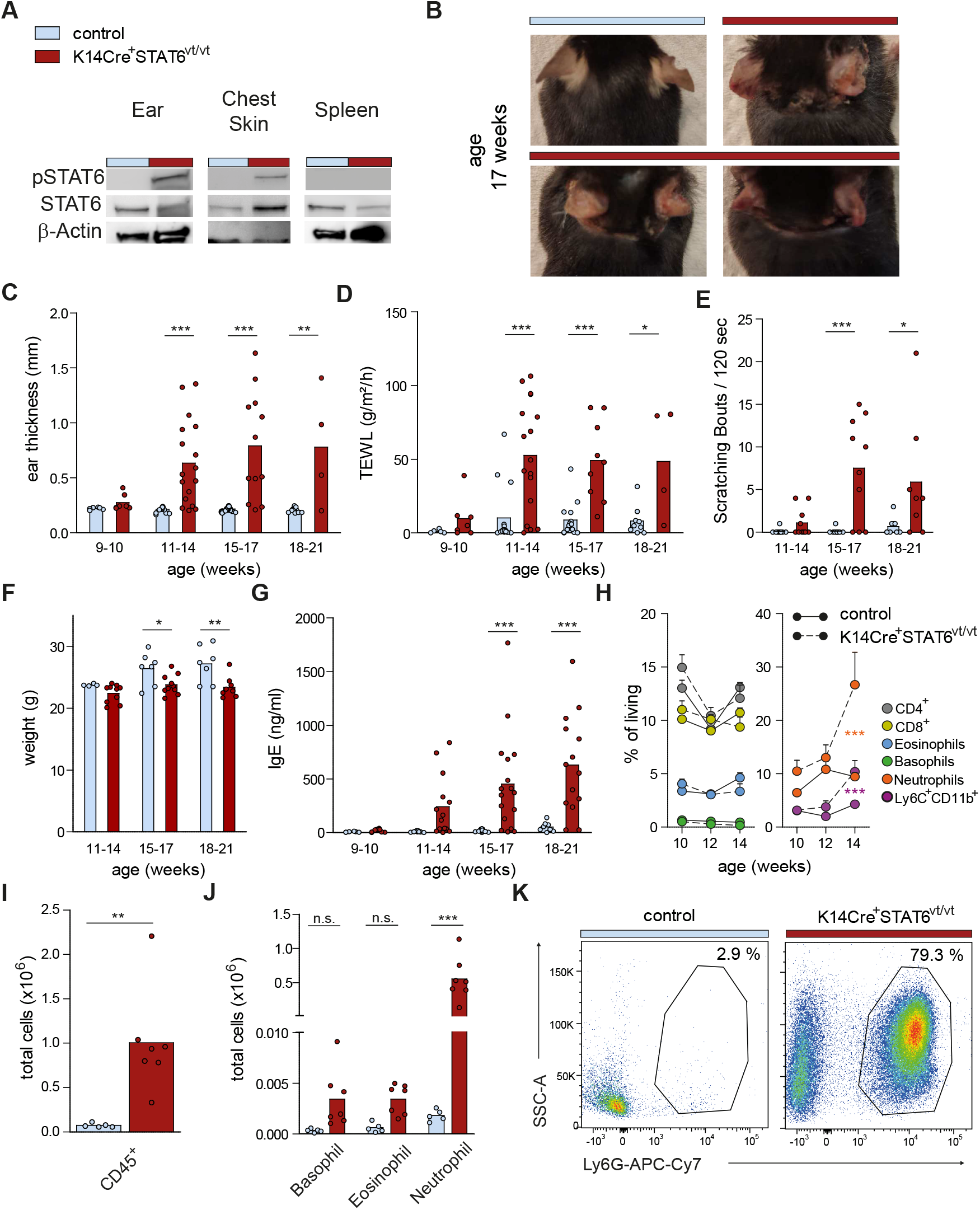
K14Cre^+^STAT6^vt/vt^ mice spontaneously develop skin lesion with underlying neutrophilia correlated with itch. In these experiments K14Cre^+^STAT6^vt/vt^ and control mice were compared. A) STAT6, phospho-STAT6 and β-Actin Western blots from skin and spleen lysates of indicated mice. B) Representative images of the neck region. C) Ear thickness and D) transepidermal water loss over time. E) Scratching behavior quantified at different age as bouts per time. F) Body weight over time. G) IgE titers in serum were determined by ELISA at indicated timepoints. H) Flow cytometric measurement of blood cells determining the percentage of CD4^+^ and CD8^+^ T cells, and Ly6C^+^CD11b^+^ cells as well as eosinophils, neutrophils and basophils. I) Ear tissue was digested and single cells were counted tocalculate the total cell number of CD45^+^ cells per ear. J) Total number of granulocytes per ear as determined by counting and flow cytometry. K) Representative flow cytometry plots of Ly6G^+^ cells (neutrophils) in ears of indicated mice. Statistical analysis: C-H and J) ANOVA with Bonferroni post-hoc test; I) Student’s t-test. Sample sizes and repetitions (n): A: Representative of two independent experiments. C & D: Four independent experiments: weeks 9-10 (6 control vs 7 K14Cre^+^STAT6^vt/vt^); weeks 11-14 (16 control vs 17K14Cre^+^STAT6^vt/vt^); weeks 15-17 (14 control vs 9 K14Cre^+^STAT6^vt/vt^); weeks 18-21 (9 controlvs 4 K14Cre^+^STAT6^vt/vt^). E: Two independent experiments: weeks 11-14 (10 control vs 10K14Cre^+^STAT6^vt/vt^); weeks 15-17 (10 control vs 10 K14Cre^+^STAT6^vt/vt^); weeks 18-21 (9 control vs 8 K14Cre^+^STAT6^vt/vt^). F: Two independent experiments: weeks 11-14 (4 control vs 10 K14Cre^+^STAT6^vt/vt^); weeks 15-17 (7 control vs 10 K14Cre^+^STAT6^vt/vt^); weeks 18-21 (7 control vs 8 K14Cre^+^STAT6^vt/vt^). G: Four independent experiments: weeks 9-10 (6 control vs 8 K14Cre^+^STAT6^vt/vt^); weeks 11-14 (9 control vs 14 K14Cre^+^STAT6^vt/vt^); weeks 15-17 (15control vs 17 K14Cre^+^STAT6^vt/vt^); weeks 18-21 (8 control vs 14 K14Cre^+^STAT6^vt/vt^). H: Two independent experiments (6 control vs 7 K14Cre^+^STAT6^vt/vt^); Error bars represent SEM. I & J: Two independent experiments (5 control vs 7 K14Cre^+^STAT6^vt/vt^).

**Figure 2.**
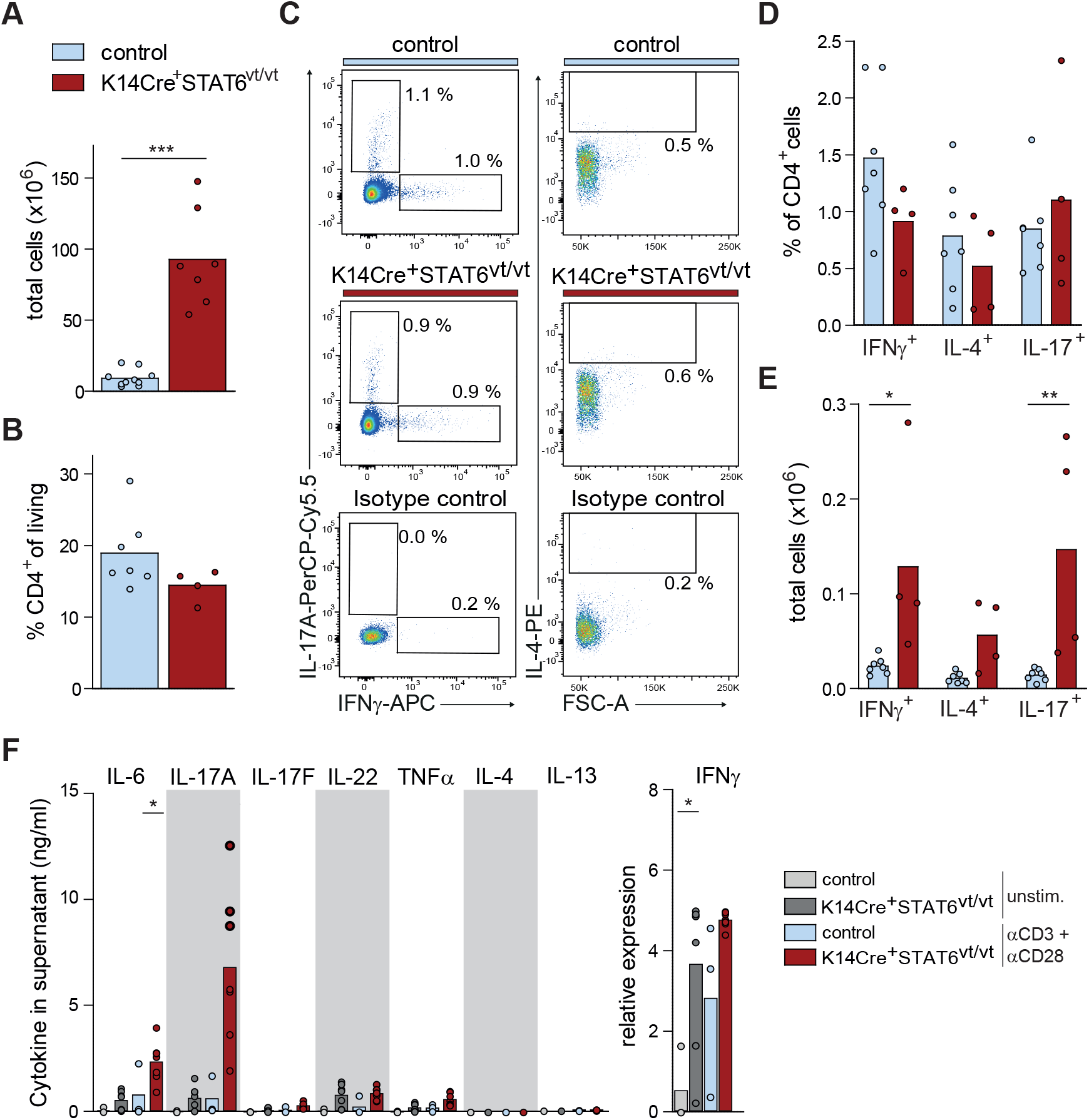
Lymph node hyperplasia and increased type 1 and type 3 cytokines in lymph nodes of K14Cre^+^STAT6^vt/vt^ mice with skin inflammation. Auricular and superficial cervical ear-draining lymph nodes (ear LNs) of K14Cre^+^STAT6^vt/vt^ and control mice were analyzed. A) Total cell count in lymph nodes. B) Percentage of CD4^+^ T cells as determined by flow cytometry. C) LN cells were restimulated with PMA/Ionomycin and intracellularly stained for IL-17, IFNγ and IL-4 and their percentage within total CD4^+^ T cells D) as well as their total cell numbers E) were determined. Statistical analysis: A-B) Student’s t-test; D-F) ANOVA with Bonferroni post-hoc test. All experiments were repeated twice. Sample sizes (n): A: 10 control vs 7 K14Cre^+^STAT6^vt/vt^. B and D-E: 7 control vs 4 K14Cre^+^STAT6^vt/vt^. F: 3 control vs 7 K14Cre^+^STAT6^vt/vt^.

Similar to inflamed ear tissue, we observed increased total cell numbers in ear-draining lymph nodes (LN) (**Fig. 2A**), but the relative abundance of CD4^+^ T cells remained unchanged (**Fig. 2B**). The proportions of Th1, Th2, or Th17 subsets among CD4^+^ T cells were also unaltered (**Fig. 2C-D, gating in Suppl. Fig. 2C**). However, considering the increased total cell number, there was a significant rise in total Th1 and Th17 cells, and a trend toward increased Th2 cells (**Fig. 2E**), potentially contributing to pathology in K14Cre^+^STAT6^vt/vt^ mice. Supporting this notion, *ex vivo* analysis of LN T cells from these mice showed enhanced baseline production of type 1 immunity-associated IFNγ and increased IL-17A (but not IL-17F) expression after T cell-specific stimulation with anti-CD3/anti-CD28. In contrast, IL-22, TNFα, or type 2 immunity-associated cytokines such as IL-4 and IL-13 were minimally produced *ex vivo* (**Fig. 2F**). These findings suggest that T cells in K14Cre^+^STAT6^vt/vt^ mice are primed to favor type 1/type 3 immune responses rather than type 2 immunity.

### STAT6-driven skin lesions are linked to *S. aureus* colonization and can be prevented by antibiotics

To study lesional and non-lesional skin from the same K14Cre^+^STAT6^vt/vt^ mouse, as well as tissue from control animals, we sampled back-/neck-skin as outlined (**Fig. 3A**). Lesional skin exhibited pronounced histopathology, including thickening of dermal layers and cell infiltration (**Fig. 3B**). Neutrophils constituted an increased proportion of total CD45^+^ cells in lesional samples, similar to inflamed ear-skin but less pronounced. In contrast, the proportions of basophils, eosinophils, mast cells, ILC2s, or CD4^+^ T cells remain unchanged, with only minor differences observed for B cells (**Suppl. Fig. 3A-B**). Analysis of skin lysates revealed elevated expression of the inflammatory cytokine IL-6 and the chemokine CCL3, which is known to recruit neutrophils^38^, in lesional skin (**Fig. 3C-D**). Given that skin diseases and neutrophilia are often associated with altered microbiomes, we performed 16S rRNA gene sequencing to determine whether STAT6 activity in keratinocytes also impacts the skin microbiome. In non-lesional skin of K14Cre^+^STAT6^vt/vt^ mice, alpha-diversity was reduced compared to wild-type controls, and this reduction was even more pronounced in lesional skin (**Fig. 3E**). Furthermore, PERMANOVA analysis revealed significant differences in the microbiome composition between all groups: wild-type, non-lesional and lesional K14Cre^+^STAT6^vt/vt^ samples (**Fig. 3F**). These differences were driven by a massive outgrowth of bacteria belonging to the genus *Staphylococcus* in both K14Cre^+^STAT6^vt/vt^-derived groups (**Fig. 3G**). qPCR analysis of the *femB* gene - a characteristic marker of *S. aureus* - confirmed this outgrowth was indeed dominated by *S. aureus* (**Fig. 3H**). To investigate whether these microbiome shifts contributed to the pathology observed in K14Cre^+^STAT6^vt/vt^ mice, a group of these mice was treated with antibiotics via drinking water to reduce microbial load. In addition, we distinguished between co-housed and non-co-housed control animals in separate cages. Skin swabs were taken over time and cultured on blood agar plates to monitor the presence of *S. aureus*. At 10 weeks of age, no significant outgrowth of *S. aureus* was observed in any group (**Fig. 3I**). However, by 17 weeks of age, *S. aureus* outgrowth was evident only in untreated K14Cre^+^STAT6^vt/vt^ mice, correlating with the presence of skin lesions. Antibiotic treatment therefore effectively prevented *S. aureus* outgrowth. Notably, co-housing of *S. aureus*-colonized K14Cre^+^STAT6^vt/vt^ mice with control mice was insufficient to drive appreciable *S. aureus* colonization in control mice. We discontinued antibiotic treatment at this time point and by 23 weeks of age, the previously treated K14Cre^+^STAT6^vt/vt^ group also exhibited *S. aureus* outgrowth (**Fig. 3I**). *S. aureus* outgrowth correlated with skin inflammation (ear swelling), underscoring the critical role of the microbiome in inducing pathology in K14Cre^+^STAT6^vt/vt^ mice (**Fig. 3J**). In summary, persistent STAT6 activity in keratinocytes is sufficient to promote *S. aureus* outgrowth on the skin, and treatment with antibiotics can prevent skin lesions in K14Cre^+^STAT6^vt/vt^ mice.

**Figure 3.**
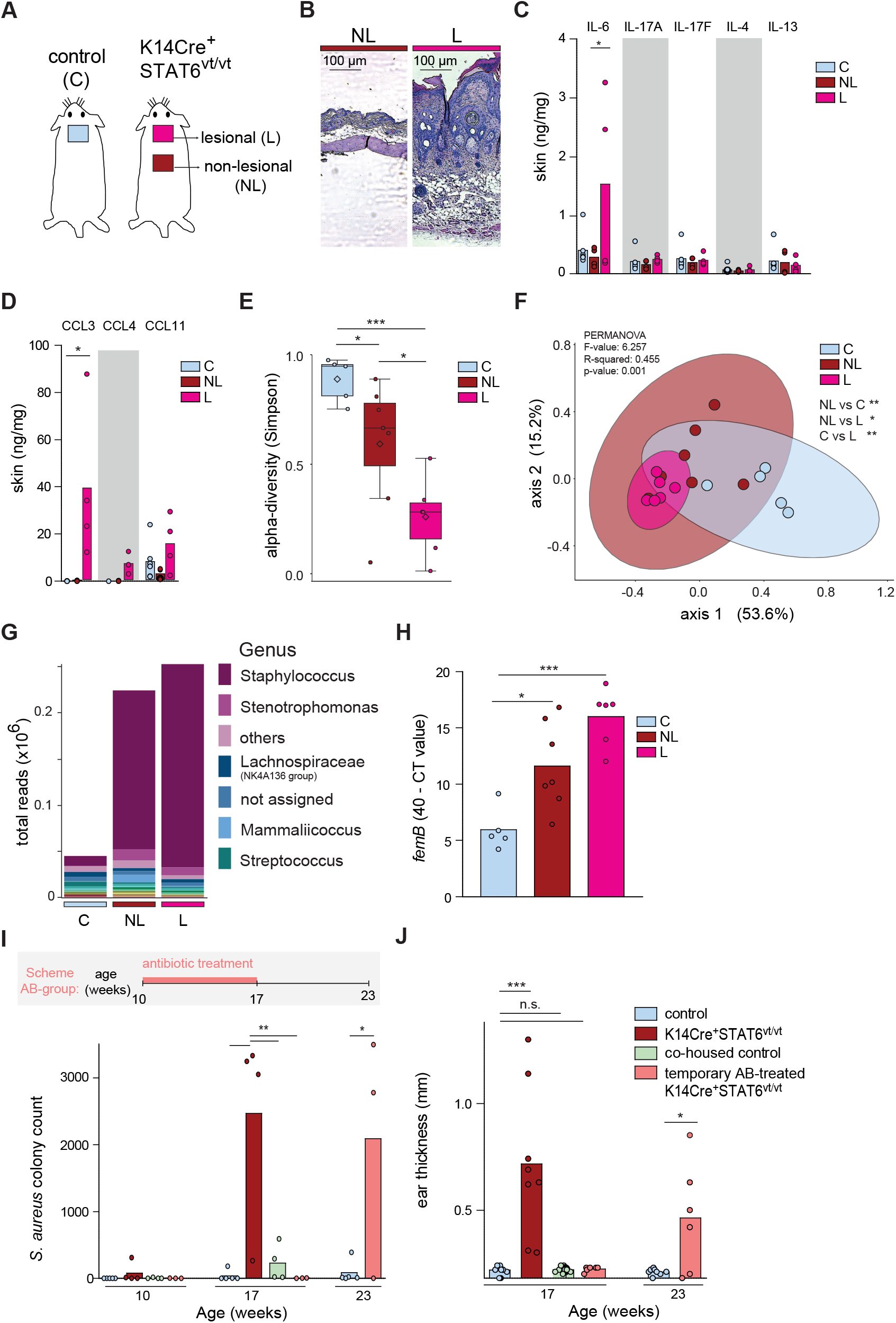
STAT6-driven skin lesions are linked to *S. aureus* expansion and can be prevented by antibiotic treatment. Comparison of back skin pathology and microbiome of K14Cre^+^STAT6^vt/vt^ and control mice in response to antibiotic treatment. A) Scheme of skin sampling sites. B) H&E staining of back skin sections. C) Cytokine and D) chemokine quantification in skin lysates. E) Alpha diversity of skin bacteria. F) Dimensional reduction visualization of differences in 16S rRNA sequencing data between groups and corresponding PERMANOVA statistics. G) Total read numbers in respective groups and subgrouping indicates genus of microbes. H) *S. aureus*-specific *femB* qPCR. I-J) Antibiotic treatment (AB) of indicated groups between age 10 - 17 weeks and determination of *S. aureus* colonization. In addition, ear thickness data over time is provided as readout of inflammation. Statisticalanalysis: C-E, H-J) ANOVA with Bonferroni post-hoc test. All experiments were repeated twice; J three times. Sample sizes (n): B: Representative images. C-D: 7 control vs 4 non-lesional vs 4 lesional. E-G: 5 control vs 7 non-lesional vs 6 lesional. H: 5 control vs 7 non-lesional vs 6 lesional. I: 5 control vs 4 K14Cre^+^STAT6^vt/vt^ vs 4 co-housed vs 3 AB group. H: Ears: 8 control vs 8 K14Cre^+^STAT6^vt/vt^ vs 8 co-housed vs 6 AB.

### Persistent STAT6-activity in keratinocytes induces a stress/proliferation gene signature and activates type 1/type 3 immunity in the skin

To gain a broader understanding of pathology-associated transcriptional changes, we performed bulk RNA sequencing on skin samples from K14Cre^+^STAT6^vt/vt^ and control mice. In non-inflamed skin, the gene expression profiles were similar between the two groups. However, the transcriptome was clearly altered in inflamed skin (**Fig. 4A-C**). Among the upregulated genes in lesional skin were anti-microbial genes (*Lce* and *Sprr*), as well as genes involved in tissue damage regulation, such as genes belonging to the Stfa family. Cytokine signature analysis revealed enrichment of IL-4/IL-13-regulated pathways in non-lesional K14Cre^+^STAT6^vt/vt^ skin compared to skin of control animals, indicating a low but measurable baseline STAT6 activity in the skin of mice that express constitutively active STAT6vt in keratinocytes (**Fig. 4D**). In lesional skin, the top upregulated gene sets were associated with type 1 immunity (IL-15, IL-12, IFNε, IL-7) and type 3 immunity (IL-36A), consistent with observed neutrophilia and *S. aureus* outgrowth in K14Cre^+^STAT6^vt/vt^ mice (**Fig. 4D**). We further checked for enrichment of murine hallmark gene sets^39^ which showed enrichment of interferon-associated genes, pathways related to *Tnf* and *Il6* signaling, and genes linked to cell activation, proliferation, metabolism, and effector functions (gene sets: E2F, G2 to M transition, Myc, mTORC1, glycolysis, oxidative phosphorylation, cholesterol homeostasis, respectively). Stress and tissue remodeling-related gene sets, such as apoptosis, DNA repair, and angiogenesis, were also upregulated in lesional skin (**Fig. 4E**). We also observed upregulation of neutrophil-associated genes upregulated, such as *Stefins* or anti-microbial *S100a8 and S100a9* which are prominent in neutrophils but can also be released by other cells including keratinocytes (**Fig. 4F**). Regarding skin barrier changes, genes associated with keratinocyte hyperproliferation and stress (*Krt6a, Krt6b, Krt14, Krt16, Krt17, Krt31, Krt36*) were upregulated, while genes linked to late differentiation stages (*Lor, Krt10, Krt24, Krt77, Krt78*) and stem cell quiescence (*Krt15*) were downregulated in lesional skin. Notably, *Flg* expression, which is a hallmark of atopic dermatitis in adults, was not significantly different among the three groups, suggesting that lesional skin in K14Cre^+^STAT6^vt/vt^ mice may rather resemble AD in children^6^ (**Fig. 4G**). Further, cytokine-, chemokine-, and itching/scratching-related genes were upregulated. While lesional skin in K14Cre^+^STAT6^vt/vt^ did not show an obvious type 2 immunity *(Il4, Il5, Il13*) or type 3 immunity (*Il17a, Il17f*) profile, genes encoding cytokines associated with keratinocyte hyperproliferation (*Il19, Il20*) or inflammation (*Il1b, Il6, Il33, Il24*) were upregulated. In accordance with the observed scratching phenotype K14Cre^+^STAT6^vt/vt^ mice, we also found upregulation of *Postn, Il1b* and *Il6*. We further found higher expression of IL-36 family genes that were reported to drive IgE production in presence of *S. aureus* ^40^ (**Fig. 4H**). Taken together, these data indicate a stress/hyperproliferative and less differentiated state of keratinocytes in lesional skin of K14Cre^+^STAT6^vt/vt^ mice. Lesional skin further shows evidence of type 1 and type 3 immunity activation, along with signs of immune cell activation, including neutrophils. In non-lesional skin, we observed no striking differences between control and K14Cre^+^STAT6^vt/vt^ mice.

**Figure 4.**
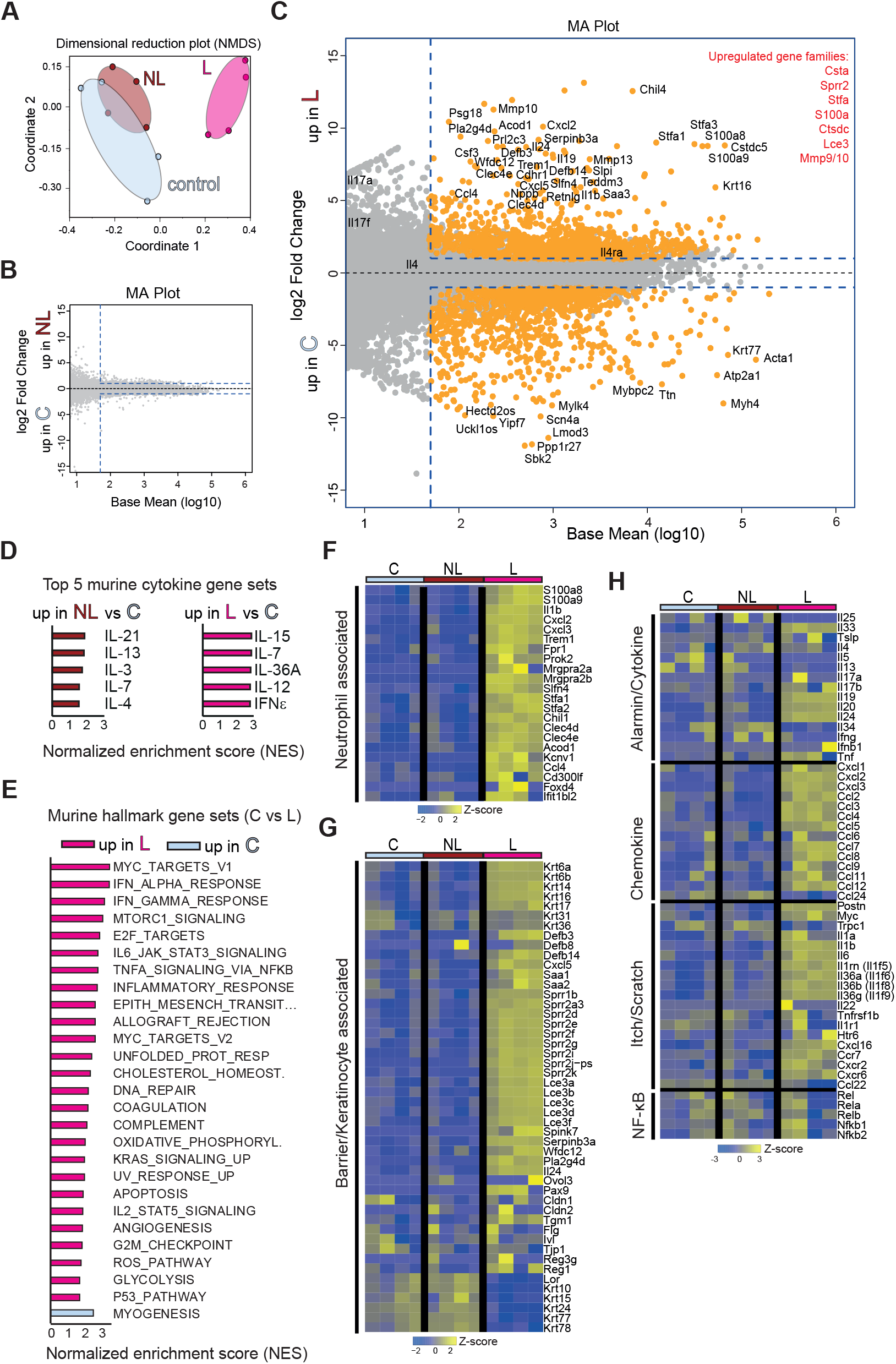
STAT6-activity in keratinocytes induces a stress/proliferation gene-signature and activates type 1/type 3 immunity in the skin. Bulk RNA sequencing of skin lysates of K14Cre^+^STAT6^vt/vt^ and control mice was performed. A) Dimensional reduction plot to visualize broad transcriptomic differences between groups (C = control, K14Cre^+^STAT6^vt/vt^: NL = non lesional, L = lesional; see scheme Fig. 3A). B) MA plot comparing the transcriptional profile of control and non lesional groups or C) control and lesional skin. D) Gene set enrichment using cytokine signature gene sets for indicated groups. Shown are the five gene sets with the highest normalized enrichment score for every comparison. E) Gene set enrichment using murine hallmark gene sets. Shown are all significantly different sets between control and lesional skin. F, G, H) Heatmaps of F) neutrophil, G) barrier/keratinocyte and H) chemokine, cytokine, itch/scratch associated genes. Samples were collected in two independent experiments, but were sequenced together. Sample sizes (n): A-H four mice per group.

### K14Cre^+^STAT6^vt/vt^ mice exhibit a transcriptomic profile consistent with chronic atopic dermatitis

As a next step, we compared the transcriptomic profile of lesional skin of K14Cre^+^STAT6^vt/vt^ mice to publicly available data sets from studies of human skin diseases. To identify relevant genes, we selected the 300 most highly upregulated genes in lesional skin of K14Cre^+^STAT6^vt/vt^ mice and analyzed expression levels of their human orthologs in single-cell RNAseq data sets from skin biopsies of AD (type 2 immunity driven/mixed responses) and psoriasis (type 1 and type 3 immunity driven) patients as well as healthy donors. For this analysis, we calculated a module score^41^, integrating the expression information for the 300 orthologs, for each cell of the published human data sets. Our analysis revealed that genes from our generated murine signature (STAT6-module genes) are expressed in human keratinocytes^42^ and especially in AD and psoriasis samples, suggesting that STAT6-induced activity in human keratinocytes contributes to pathology in AD and psoriasis (**Fig. 5A**). To identify immune cell subpopulations expressing these genes, we analyzed another human single-cell RNAseq data set of CD45^+^ cells which contains high-resolution information on subpopulations^43^, comparing expression levels in skin from AD and psoriasis patients versus healthy controls. For each subpopulation, we also provide information about the percentage of cells derived from the three different disease states (AD, psoriasis, healthy) to reveal disease-specific alterations in population size. Strong signature scores were observed in dendritic cell, monocyte, and Langerhans cell clusters, suggesting a conserved role across both mouse models and human diseases. Notably, similar patterns emerged in AD and psoriasis (**Figs. 5B-C**), suggesting shared pathogenic mechanisms. As we hypothesize that STAT6 activity in keratinocytes resembles a late or chronic stage of AD, we also determined expression of our STAT6-module genes in a bulk RNAseq data set that includes chronic AD samples^44, 45^. Heatmap visualization showed that the STAT6-module genes are upregulated in AD samples, especially chronic AD supporting the notion that the pathology in K14Cre^+^STAT6^vt/vt^ mice resembles a chronic stage of AD. However, STAT6-module genes were also upregulated in psoriatic samples, highlighting shared features of chronic AD and psoriasis, such as their association with type 1 and type 3 immunity (**Fig. 5D**). Principal component analysis on the human bulk RNAseq data set, based on STAT6-module genes, revealed that the first two components are sufficient to separate healthy, AD, and psoriasis samples. The cluster of chronic AD samples is partly separated from the acute AD samples, and the gene eigenvalues indicate that *KRT6A* (stress induced), *SERPINB4* (alarmin), *S100A8* and *S100A9* (alarmins mainly produced by neutrophils or activated keratinocytes) drive this separation. It also points to a relevant role of neutrophils in chronic AD, that might be often missed in single cell datasets due to technical challenges in single-cell RNAseq analyses of granulocytes. In summary, these data suggests that orthologs of genes expressed in lesions of K14Cre^+^STAT6^vt/vt^ mice are also relevant to drive chronic AD.

**Figure 5.**
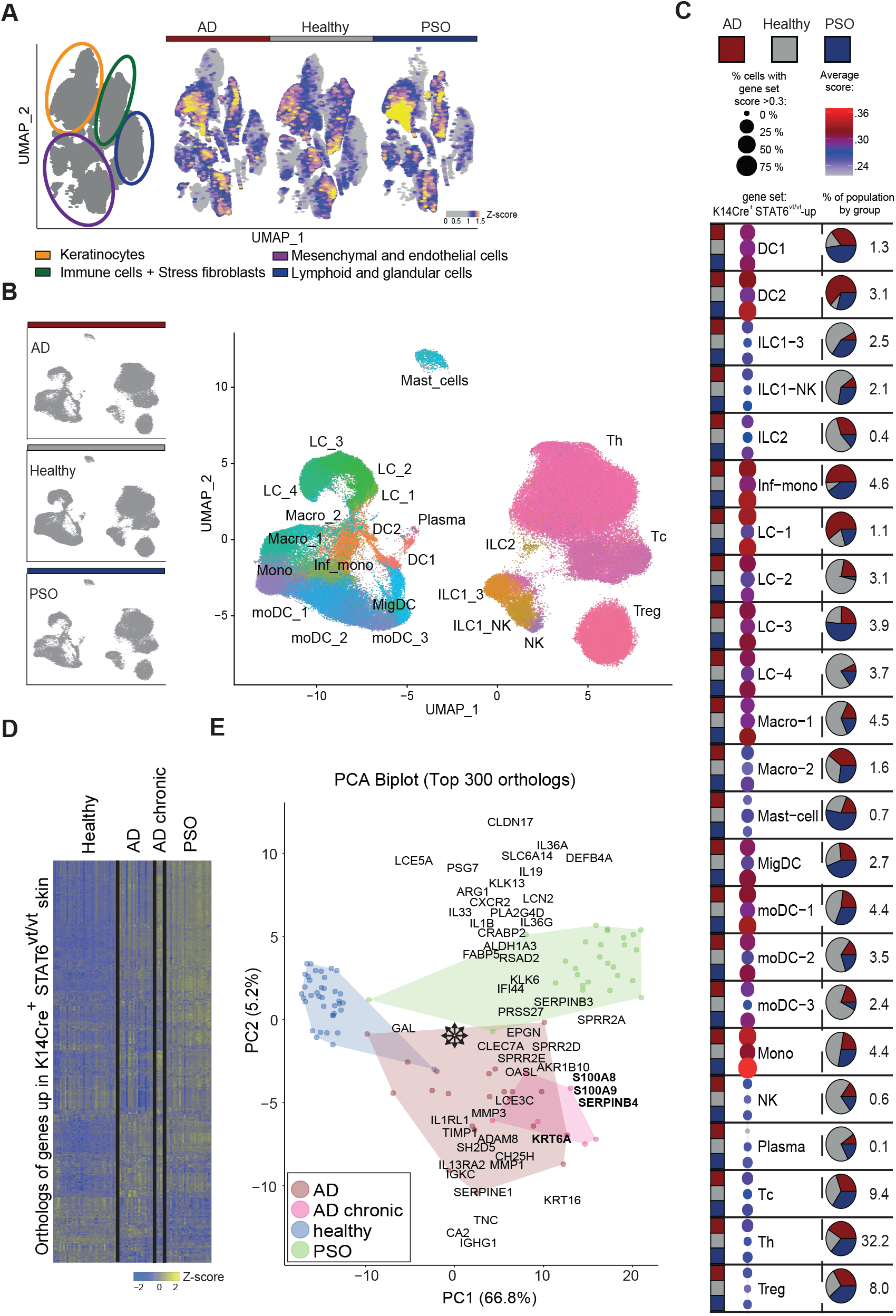
Transcriptional changes in K14Cre^+^STAT6^vt/vt^ mice resemble profiles of human skin-disease. A gene set comprising human orthologs of the 300 most upregulated murine genes in lesional skin from K14Cre^+^STAT6^vt/vt^ mice, compared to healthy skin, was generated and expression in human datasets was analyzed. A) A gene set score based on the murine lesional gene set was calculated for a published single cell dataset including keratinocytes of healthy skin, AD and psoriasis. The score is indicated by the color of the dots in the UMAP. B) Murine lesional gene set score for single cell RNAseq data of CD45^+^ cells from healthy or diseased human skin. C) A gene set score for every cluster and disease type of the CD45^+^ cells is visualized and the proportion of cells from the conditions healthy, AD and psoriasis are depicted. D) The human orthologs from the murine lesional skin gene set (STAT6-module genes) were selected from human bulk RNAseq data and were plotted for healthy skin, AD, chronic AD and psoriasis. E) Dimensional reduction of data using PCA. Each point represents an individual. On top, selected genes are overlaid. The direction of the gene name seen from the zero point shows in which direction and with what relative strength the expression of the gene shifts the samples. This gives a visual impression, showing which genes drive sample separation in the PCA plot. Sample sizes (n): A: Samples: 5 control vs 4AD vs 4 Psoriasis. B-C: Samples: 7 control vs 7 AD vs 8 Psoriasis. D-E: Samples: 38 control vs 21 AD vs 28 Psoriasis vs 6 chronic AD.

### In a type 2 immunity-driven mouse model of AD, persistent STAT6 activity in keratinocytes induces a mixed immune response resembling chronic AD in humans

Initially, type 2 immunity dominates AD pathogenesis before transitioning to mixed inflammation during chronification. However, lesional skin of K14Cre^+^STAT6^vt/vt^ does not exhibit a pronounced type 2 immune signature but rather displays features of type 1/type 3 immunity. To further investigate this type 1-/type 3-biased inflammatory response, we utilized an inducible TSLP-driven AD model based on topical application of the vitamin D3 analog MC903 (calcipotriol) to the ear skin. Thereby, we could assess if it also results in a mixed response, as often observed in the transition to chronic AD in human patients. Instead of using K14Cre^+^STAT6^vt/vt^ homozygous mice, we utilized K14Cre^+^STAT6^vt/wt^ heterozygous mice, which are less prone to developing spontaneous skin inflammation, allowing us to investigate MC903-induced inflammation. We first confirmed that STAT6 is constitutively phosphorylated in the skin of heterozygous K14Cre^+^STAT6^vt/wt^ mice (**Fig. 6A**). Upon MC903 treatment, K14Cre^+^STAT6^vt/wt^ mice demonstrated a more pronounced ear swelling response as compared to control mice, consistent with the severe disease observed in chronic AD patients (**Fig. 6B-C**) along with reduced *Il4* expression (**Fig. 6D**). This was associated with a mixed granulocyte response characterized by eosinophils and neutrophils. Notably, the control group showed an increase of eosinophils but not neutrophils (**Fig. 6E**). This demonstrates that persistent STAT6 activity in keratinocytes promotes a mixed immune response with neutrophilia, even in an otherwise eosinophil-associated AD model, thereby reflecting the situation frequently seen in chronic AD patients.

**Figure 6.**
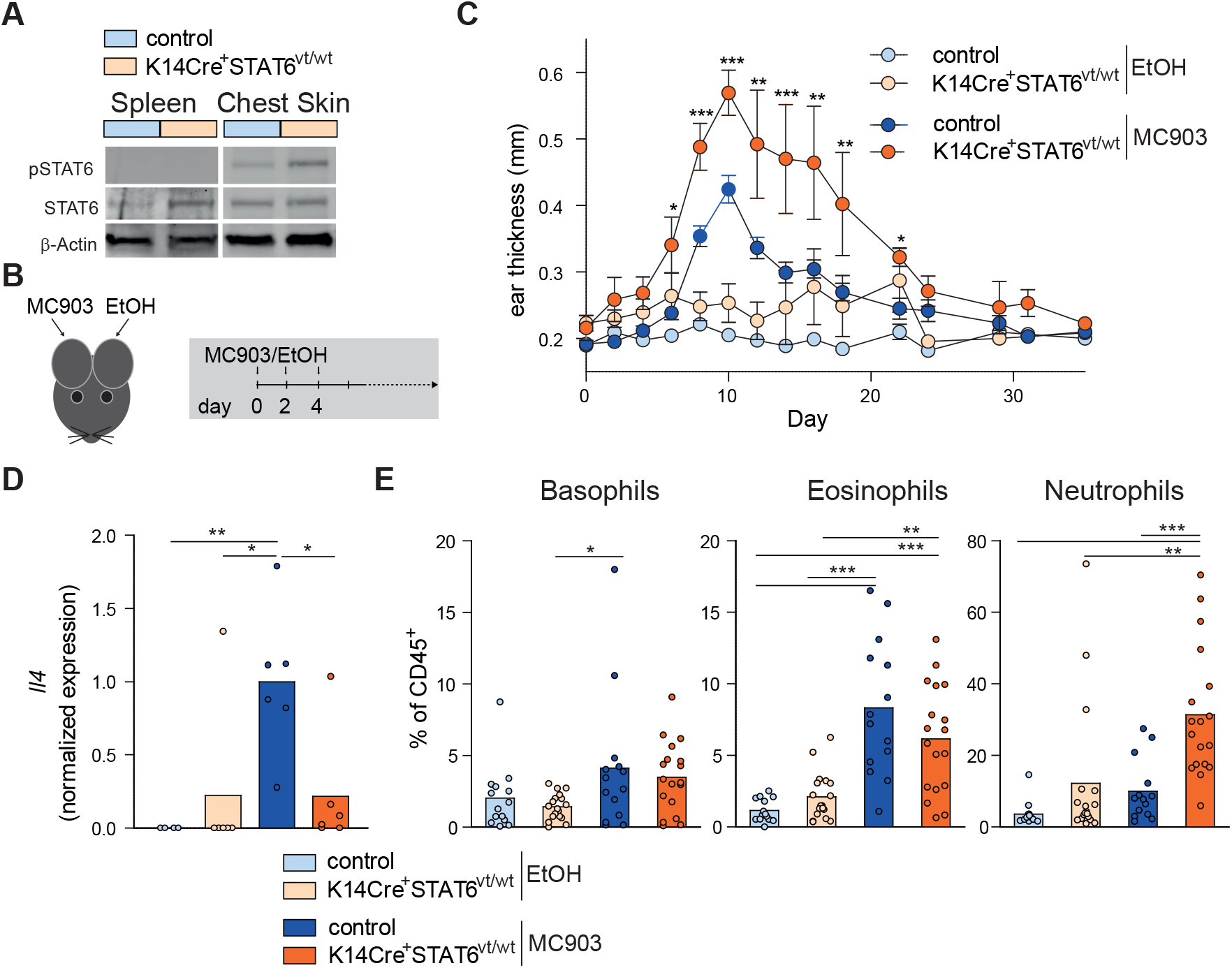
MC903-induced AD-like ear inflammation is stronger, prolonged and accompanied by neutrophilia in K14Cre^+^STAT6^vt/wt^ mice. Heterozygous K14Cre+STAT6vt/wt mice and littermate controls were used for all experiments. A) STAT6, phospho-STAT6 and β-Actin Western blots from skin and spleen lysates as indicated. B) Treatment scheme to induce AD-like inflammation. 4 nmol of the vitamin D3 analog MC903 (calcipotriol) in ethanol or ethanol alone (EtOH) were applied on respective ears at each treatment timepoint. C) Ear thickness measurement for indicated groups over time. D) *Il4* qRT-PCR of ears from indicated groups at treatment day 10. E) Percent of basophils, eosinophils and neutrophils of CD45^+^ cells determined by flow cytometry at treatment day 10. Statistical analysis: D-E) ANOVA with Bonferroni post-hoc test. Sample sizes and repetitions (n): A: Representative of two experiments. C: Four independent experiments; days 0-10: all groups ≥ 16; days 12-18: all groups ≥ 9; days 20-22: all groups ≥ 4; days 24-35: all groups ≥ 3; Error bars represent SEM. D: All groups n=6; two independent experiments. E: Ears: 14 controls with EtOH vs 18 K14Cre^+^STAT6^vt/wt^ with EtOH vs 14 controls with MC903 vs 18 K14Cre^+^STAT6^vt/wt^ with MC903; four independent experiments.

## Discussion

In this study, we employed a new genetic approach to specifically analyze the outcome of persistent STAT6 activity in keratinocytes *in vivo*. Previous work has largely investigated the keratinocyte-specific impact of type 2 immunity-associated cytokines IL-4 and IL-13 and their STAT6-dependent change of the transcriptome *in vitro*, limiting conclusions about effects on clinical disease^20, 46^. Furthermore, mouse models and skin biopsies from human AD patients demonstrated an IL-4/IL-13-driven impairment of skin barrier integrity likely resulting from reduced STAT6-driven downregulation of tight junction and extracellular matrix proteins in keratinocytes ^19, 23, 47, 48^. Here, we observed spontaneous skin lesions characterized by neutrophil infiltrates in homozygous K14Cre^+^STAT6^vt/vt^ mice. In human, the role of neutrophils in atopic dermatitis (AD) remains incompletely understood, but research in this area is rapidly expanding^49^. Skin-neutrophilia appears more pronounced in intrinsic AD compared to the extrinsic form, and is more frequently observed in children and in the Asian form of AD ^4, 5, 7, 49^. The resemblance of the murine phenotype to pediatric AD is further supported by the absence of alterations in *Flg* expression^5^ and the lack of Th22 activity^50^. As such, our model differs from previous observations of neutrophilia in *Flg* mutant mice^51^. We further noticed increased expression of IL-24, which stimulates acanthosis, scratching, and the release of neutrophil-recruiting chemokines^52, 53^. Neutrophils can promote itching in a CXCR3, CXCL1/CXCL10-dependent manner, and their ablation alleviates itch^54^, suggesting that neutrophils likely contribute to the ‘itch-scratch cycle’ observed in K14Cre^+^STAT6^vt/vt^ mice. However, this cycle can be fueled by various factors, including *S. aureus* colonization, which has been demonstrated in mice^55^ and suggested in human patients^56^. Notably, we also observe *S. aureus* colonization on both lesional and non-lesional skin of K14Cre^+^STAT6^vt/vt^ mice, mirroring the situation in human AD^57^. Antibiotic treatment prevented disease development in our mouse model, and might help to reduce infection-induced type 1/type 3 immunity features of human AD. *S. aureus* skin colonization can be increased by neutrophils through induction of oxidative stress^58^, and enhanced colonization is associated with increased itch^59^, further driving the ‘itch-scratch cycle’ and increasing disease pathology. Interestingly, we observed a relatively low level of type 2 immunity within the skin lesions of K14Cre^+^STAT6^vt/vt^ mice, indicating that STAT6-regulated genes in keratinocyte rather promote inflammation characterized by type 1 and type 3 immune responses reflective of chronic AD. This observation parallels findings in human lesions, where IL-4Rα and IL-4 levels are also decreased in chronic AD^60^. While we observe increased IgE levels, which may be mediated by *S. aureus*-induced IL-36 expression^40^, this does not appear to be a crucial driver of pathology, as IgE-deficient bone marrow-chimeric K14Cre^+^STAT6^vt/vt^ mice still developed spontaneous inflammation and pathology. This is in line with reports showing limited efficacy of human anti-IgE treatment in AD^61, 62^. Although induced IgE may contribute to atopic comorbidities^63^, it does not appear to be essential for AD pathogenesis. It has been described that type 2 immunity can impede type 1 immunity-dependent clearance of bacterial infections^64^; however, the relatively low level of type 2 immunity and the stronger type 1 immunity observed in K14Cre^+^STAT6^vt/vt^ mice do not seem sufficient to clear *S. aureus* infection or prevent disease progression. Despite the less pronounced type 2 immunity in chronic AD lesions, anti-IL-4/IL-13 receptor treatment (dupilumab) remains effective^65^. Our findings suggest that dupilumab treatment, which acts upstream of STAT6 activation in keratinocytes, provides a potential explanation for the efficacy of dupilumab even in treating chronic AD with mixed immune responses. As such, cell-type specific inhibition of STAT6 might enable more precise treatment in the future. Evolutionarily, it is tempting to speculate about the shift from type 2 to type 1/type 3 immunity. It may reflect a failure in immunity induction and resolution, or represent a necessary adaptation to prevent unsuccessful type 2 responses with barrier impairment. Our findings highlight the utility of K14Cre^+^STAT6^vt/vt^ mice as a new model for studying chronic AD, particularly in exploring phase- or endotype-specific questions. This model provides insights into disease mechanisms and potential therapeutic targets, paving the way for more effective treatments tailored to specific immune profiles.

## Material and Methods

### Mice

K14Cre^+^STAT6^vt/vt^ mice were generated by crossing mice encoding a constitutively active form of STAT6 in the Rosa26 locus behind a floxed stop cassette^66^ to mice expressing the Cre recombinase under control of the murine keratin-14 (K14) promoter^32^ which is predominantly active in basal epidermal keratinocytes. Littermates were used as controls. Bone marrow of IgE-deficient (IgEKO) mice^37^ was used to generate chimeras. All mice were on a C57BL/6 background. Animal experiments were approved by the Government of Lower Franconia and performed in accordance with the German animal protection law.

### Determination of scratching bouts per time

Mice were monitored for 120 seconds and bouts were counted. We counted one bout as the process starting from the moment an animal lifted its leg to scratch until it put it down again.

### T cell restimulation

Single-cell suspensions of lymph node cells (1×10^6^ cells/well) were restimulated in RPMI medium (10% FCS, 2 mM L-glutamine, 10 µg/mL streptomycin, 100 IU/mL penicillin) supplemented with 10 ng/mL IL-2 in plates that had been pre-coated overnight at 4°C with 1 µg/mL anti-CD3 (clone 145-2C11) and anti-CD28 (clone 37.51) antibodies (both Invitrogen) in PBS. Cells were incubated for 24 hours at 37°C with 5% CO_2_, and supernatant was collected for ELISA.

### Western Blot

Skin tissue lysates were probed for pSTAT6 (clone D8S9Y), total STAT6 (polyclonal), and β-actin (clone 13E5) using rabbit primary antibodies (Cell Signaling Technology). Following overnight 4°C incubation, HRP-conjugated donkey anti-rabbit secondary antibody (Cell Signaling Technology) and Signal Fire ECL substrate were used for detection via a Bio-Rad ChemiDoc Imaging System.

### RNA Isolation and qPCR

A part of skin (∼1cm x 1cm) was put in a 2ml ruptor tube containing ceramic Beads (1.4 mm and 2.8 mm bead mix) with RTL buffer from the RNeasy Mini Kit (Qiagen). The tissue was homogenized with OMNI Bead Ruptor (Biolabproducts GmbH, Bebensee, Germany) (3 × 5 m/s each 15 s with cooling breaks on ice). Supernatant was used for further RNA isolation. Reverse transcription of RNA was performed with the High-Capacity cDNA Reverse Transcription Kit (Thermo Fisher Scientific) according to the manufacturer’s instructions. Quantitative RT-PCR was performed using SYBR Select Master Mix (Thermo Fisher Scientific) and the following primers: Hprt-fwd 5′-GTTGGATACAGGCCAGACTTTGTT-3′, Hprt-rev 5′-GAGGGTAGGCTGGCCTATAGGCT-3′, Il4-fwd 5’ACTTGAGAGAGATCATCGGCA-3’; Il4-rev 5’-AGCTCCATGAGAACACTAGAGTT-3’. Expression of target genes was normalized to Hprt. *FemB* of *S. aureus* was detected by qPCR as previously described^67^.

### Antibiotic treatment and determination of S. aureus colonization

Drinking water supplemented with Vancomycin (0.5 mg/ml) and Ampicillin (1 mg/ml) (abbreviated as AB; both Carl Roth) was given to indicated groups between 10 and 17 weeks of age. Ear swabs were taken and spread on blood agar to detect bacterial growth. Morphologically distinct colonies were sampled and *S. aureus* was determined by clumping test and MALDI-TOF. Colonies with the same morphology were counted as *S. aureus*.

### RNAseq analysis

A part of skin (∼1 cm x 1 cm) was placed in a 2 ml ruptor tube containing ceramic Beads (1.4 mm and 2.8 mm bead mix) with RTL buffer and ß-ME from the RNeasy Mini Kit (Qiagen). The tissue was homogenized with an OMNI Bead Ruptor (Biolabproducts GmbH, Bebensee, Germany) (3 × 5 m/s each 15 s with cooling breaks on ice). Supernatant was used for further RNA isolation with the RNeasy Mini Kit according to manufacturer’s instructions. RNA was sent to Novogene Europe (Cambridge) for sequencing as a service. There, library preparation was performed with NEBNext® Ultra™ RNA Library Prep Kit for Illumina (NEB, Frankfurt, Germany) following manufacturer’s recommendations and paired-end sequencing was performed on an Illumina NovaSeq 6000. Filtered clean reads were aligned to reference genome mm10 using STAR (v2.6.1d)^68^. FeatureCounts (v1.5.0-p3) was used to generate counts used for downstream analysis in the laboratory^69^. Data were loaded into R (3.5.3; The R Foundation for statistical computing, Vienna, Austria) and genes with zero counts were removed. Deseq2 (1.20.0) was used to normalize counts and perform differential expression analysis without further shrinkage^70^. Heatmaps of log2 normalized counts were drawn with the ggplot2 package. For gene set enrichment GSEA (4.33) with normalized count values as input and recommended settings (default values but permutation type set to “gene_set”) was used^71^. fgsea (1.4.0) was used for analysis of the cytokine genesets. For the GSEA analysis selected gene sets of MSigDB database version 2025.1 for Mus musculus were used. For gene set enrichment, GSEA 4.33 software with normalized count values as input and recommended settings (default values but permutation type set to “gene_set”) was used. For hallmark gene set analysis, the MSigDB v2023.2.Mm database was employed39. Data was deposited at ArrayExpress: E-MTAB-16329. Single cell datasets from the literature were obtained via corresponding repositories (https://ftp.ebi.ac.uk/pub/databases/microarray/data/atlas/sc_experiments/E-MTAB-8142,https://zenodo.org/records/5228495, GEO: GSE121212) and analyzed using R (4.2.0), Seurat_5.0.1 and ggplot2.

### 16S rRNA sequencing

Swabs (OmniSwab, Qiagen GmbH, Hilden, Germany) moistened in sterile water were rubbed over the skin of the outer ear side 30 times. DNA was isolated from the OmniSwabs using the QIAamp UCP DNA Micro Kit according to the manufacturer’s instructions (Qiagen GmbH, Hilden, Germany) and used for 16S rRNA sequencing (Illumina). Genomic DNA was used in 30 cycle PCR amplification (NEBNext Q5 Hot Start Hifi PCR Master Mix, New England Biolabs) of genomic 16S ribosomal RNA V4 regions using the prokaryotic primer pair (515F forward primer: 5’-GTGYCAGCMGCCGCGGTAA−3’; 806R reverse primer: 5’-GGACTACNVGGGTWTCTAAT−3’) containing 12 bp barcodes on the forward primer (https://earthmicrobiome.org/protocols-and-standards/16s/). PCR products were purified with AMPure XP Beads (Beckmann Coulter), pooled in equimolar ratios and analyzed by 2 × 151 paired-end sequencing on an Illumina MiSeq device. Raw fastq files were then imported and analyzed in QIIME2 v2024.10 with DADA2 as the method for quality control, dereplication and amplicon sequence variant (ASV) table generation. MicrobiomeAnalyst 2.0 was used for downstream data analysis^72^. Data was deposited at the European Nucleotide Archive: PRJEB104316.

### Generation of bone marrow chimeras

Bone marrow from IgE-deficient (IgEKO) or wild-type donor mice was transferred into K14Cre^+^STAT6^vt/vt^ recipients that had been irradiated with 11 Gy before cell transfer. Antibiotics were applied via drinking water for four weeks and mice were analyzed at least eight weeks after reconstitution with bone marrow.

### Flow Cytometry

Single-cell suspensions were prepared from ear pinnae, ear lymph nodes, and lungs (which were PBS perfused). Lung tissue was further digested with DNase DN25 (Sigma–Aldrich) and Liberase TM (Roche, Basel, Switzerland) (both 100 µg/ml) for 30 min at 37°C with agitation. Ear pinnae were dissociated using the Multi Tissue Dissociation Kit I and gentleMACS Octo Dissociator (Miltenyi Biotec) (Fig. 1) or they were dissociated 90 minutes in media containing Liberase TM and DNAse I (as above), with continuous shaking at 37°C (Fig. 6). Spleen and lung suspensions underwent erythrocyte lysis with ACK buffer (0.15 M NH4Cl, 1 mM KHO3, 0.1 mM Na2EDTA). Lymph nodes and digested tissues were mashed through 70- or 100-μm cell strainers to obtain single cell suspensions. Then cells were incubated with Fc-Block (clone 2.4G2) and stained with fluorescent-dye conjugated antibodies for 20 minutes at 4°C. When more than one BD Horizon Brilliant reagent coupled antibody was in the stain BD Horizon Brilliant Stain Buffer was added. Used antibodies and staining reagents were anti-CD4-BV711 or BV421 (RM4-5), anti-CD49b-Af647 (HMα2), anti-Ly6G-APC-Cy7 or FITC (1A8), anti-CD11b-Af488 or BV786 (M1/70), anti-Ly6C-PE-Cy7 (HK1-4), anti-CD200R3-APC (Ba13) (all BioLegend), anti-CD8a-BV737 or BUV395 (53-6.7), anti-Siglec-F-BV421 or PE (E50-2440), anti-IgE-BB700 (R35-72), anti-CD45-PerCP-Cy5.5, anti-CD127-BV711 (SB/199), anti-Thy1.2/CD90.2-APC-Cy7 (53-2.1), anti-B220-BV496 (RA3-6B2), anti-CD3e-BUV395 (145-2C11), anti-CD11c-BUV395 (HL3), anti-CD335/NKP46-BUV395 (29A1.4), anti-NK1.1-BUV395 (PK136), anti-ST2-BV421 (U29-93) (all BD), anti-FcεR1 (MAR-1), anti-CD8a-APC-eFluor 780 (53-6.7) (eBiosciences), CD117-PE-Cy7 (2B8), Fixable viability dye eFluor 506 (all Invitrogen), and Fc receptor blocking Ab (anti-CD16/32, 2.4G2) (BioXCell). For intracellular cytokine staining, cells were restimulated with PMA (50 ng/ml) and ionomycin ionomycin (2 µg/ml) for 1.5 h, followed by GolgiPlug addition (1:1000) for another 2 h. Cells were surface stained, fixed/permeabilized using the BD Cytofix/Cytoperm Kit (BD), and then stained overnight at 4°C with anti-IL-4-PE (11B11), anti-IFNγ-APC (XMG1.2), and anti-IL-17a-PerCP-cy5.5 (TC11-18H10). Flow cytometric analysis was performed on a BD LSRFortessa and data analyzed using FlowJo software (BD).

### Ear Thickness and Transepidermal Water Loss (TEWL) Measurement

Ear thickness was measured with an electronic caliper (INSIZE, Suzhou New District, China). Tewameter TM Nano device (Courage + Khazaka Electronic GmbH, Koeln, Germany). Was used to measure transepidermal water loss (TEWL). TEWL was measured at the outer side of the ear for at least 90 seconds until values stabilized.

### ELISA

Total IgE serum levels were quantified by ELISA. Briefly, microplates were coated with anti-IgE (clone R35-72, BD), blocked with 3% BSA, and probed with IgE κ isotype standards (BD). Detection utilized alkaline phosphatase-conjugated anti-mouse IgE (SouthernBiotech), and absorbance at 405 nm was measured using a Multiskan FC Microplate Photometer (ThermoFisher Scientific) following para-nitrophenyl substrate conversion (Sigma-Aldrich).

### Cytometric Bead Assays

For bead-based cytokine and chemokine measurement, a part of skin (∼1cm x 1cm) was put in a 2 ml ruptor tube with 400 µl RIPA buffer (1% NP-40, 50 mM Tris pH 7.4, 0.15 M NaCl, 1 mM EDTA, 0.5% C24H39NaO4) supplemented with Protease Inhibitor and Phospho STOP (Roche). The tissue was homogenized with OMNI Bead Ruptor (Biolabproducts GmbH, Bebensee, Germany) (3 × 5 m/s each 15 s with cooling breaks on ice) in RIPA lysis buffer and centrifuged at 14000 rpm for 10 min. The supernatant was analyzed with LEGENDplex Kits 741068 and 740451 (BioLegend).

### Histology

Paraffin sections (10 μm thick) were generated from back skin. Sections were stained with hematoxylin and eosin and imaged with an Axio Vert. A1 microscope (Zeiss, Jena, Germany).

### Statistical analysis

Graph Pad Prism 5 (GraphPad Software, Boston, MA) was used for statistical analysis. Used Statistical tests are mentioned in figure legends. Two-tailed tests for significance were chosen. Significance is indicated as: *p ≤ 0.05, **p ≤ 0.01, ***p ≤ 0.001.

## Acknowledgement

We thank Kirstin Castiglione and Daniela Döhler for excellent technical support and members of the Voehringer lab for helpful discussions. Financial support was granted by the German Research Foundation (Deutsche Forschungsgemeinschaft, DFG) projects VO944/9-2 (322359157), VO944/13-1 (500725705), SFB1181_A02, and RTG2740_A7A (447268119) to D.V., RA3732/1-1 and RA3732/1-3 to D.R., TRR241_A03 (375876048) and WI 3304/8-1 (505539112) to S.W., and the Staedtler Stiftung to D.V..

## Author contributions

DR and DV designed experiments and wrote the manuscript; EC, LG, JP and DR performed experiments; DP and WG helped to identify S. aureus by mass-spec and PCR; SW performed 16S RNAseq; DR analyzed bulk and single-cell RNAseq data; SE provided K14Cre mice; all authors read and revised the manuscript.

## Data availability statement

All primary data are available from the corresponding author upon reasonable request.

