## Supplemental Figures for "Persistent activation of STAT6 in keratinocytes elicits neutrophilic skin inflammation, pruritus and *S. aureus* colonization reminiscent of chronic atopic dermatitis"

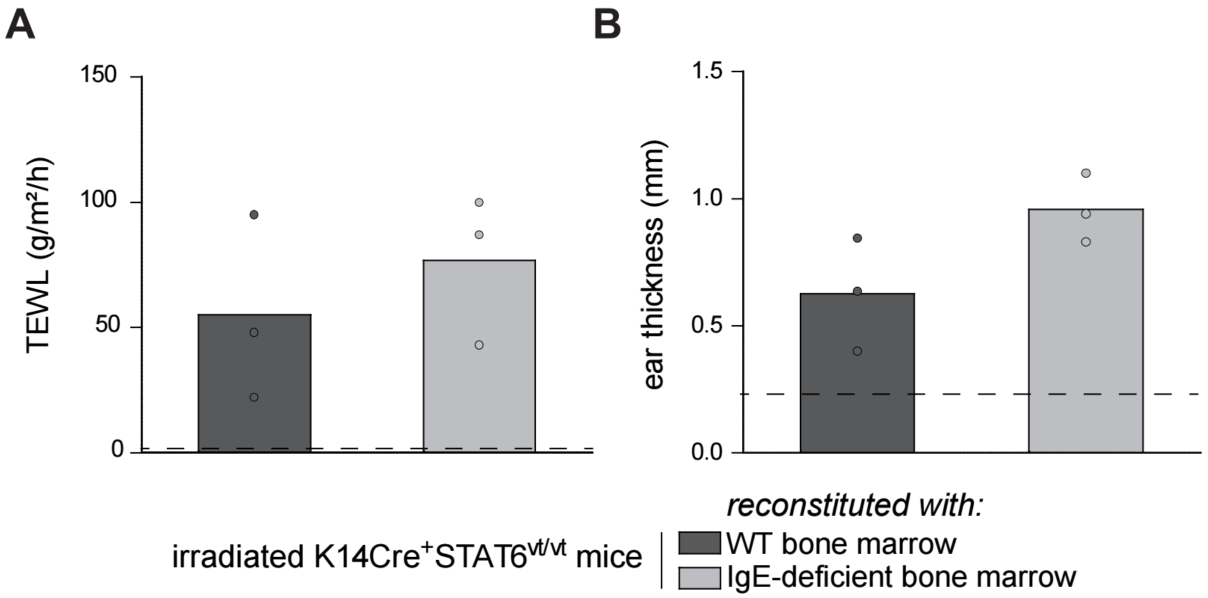


**Supplementary Figure 1. Chimeric K14Cre$^{+}$STAT6$^{vt/vt}$ mice with IgEKO hematopoietic cells spontaneously develop skin inflammation.** Analysis of irradiated K14CreSTAT6$^{vt/vt}$ mice reconstituted with IgEKO hematopoietic cells or wild-type (WT) hematopoietic cells. A) Transepidermal water loss (TEWL) and B) ear thickness. A) and B) Dashed lines indicate TEWL and ear thickness for WT control mice. Student's t-test was performed for statistical analysis and no significant differences between groups were observed.

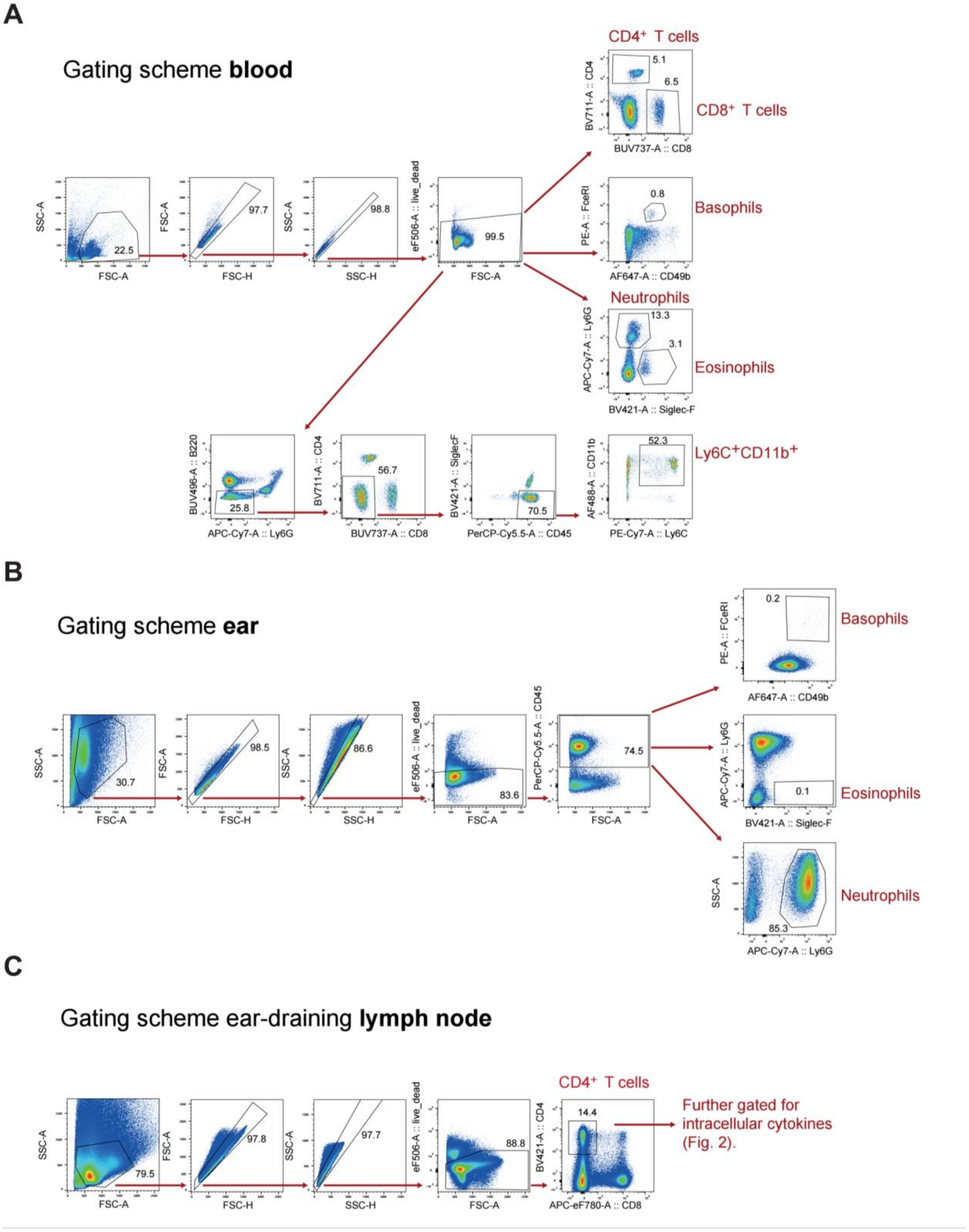


**Supplementary Figure 2. Gating schemes for flow cytometric measurements presented in Figures 1 and 2.** Gating schemes for A) blood, B) ears, and C) auricular and superficial cervical ear-draining lymph nodes.

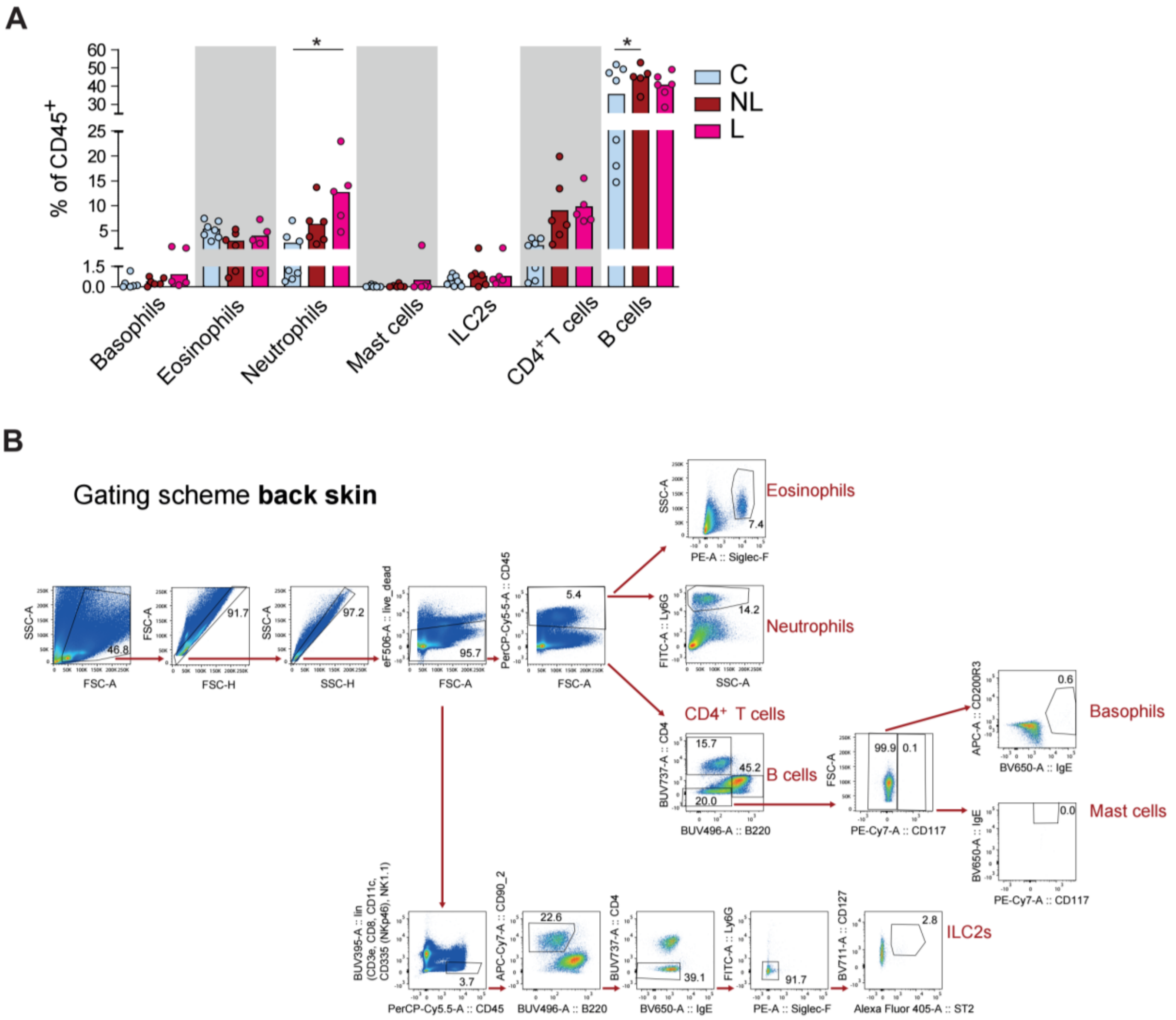


**Supplementary Figure 3. Neutrophilia is observed in lesional back skin of K14Cre$^{+}$STAT6$^{vt/vt}$ mice.** A) Flow cytometric analysis of back skin for basophils, eosinophils, neutrophils, mast cells, ILC2s, CD4$^{+}$ and B cells amongst the CD45$^{+}$ cells with B) corresponding gating scheme. Statistical analysis in A) was performed with ANOVA and Bonferroni post-hoc test. *, $p<0.05$.
